# Integrity of anterior corpus callosum is well related to language impairment after traumatic brain injury

**DOI:** 10.1101/2020.06.12.148569

**Authors:** Hae In Lee, Minjae Cho, Yoonhye Na, Yu Mi Hwang, Sung-Bom Pyun

## Abstract

**Background:** The corpus callosum (CC) serves as the bridge that relays information between the two cerebral hemispheres, and is one of the most commonly injured areas after traumatic brain injury (TBI). This study was designed to investigate the association between the CC integrity and language function after TBI.

**Methods:** We retrospectively enrolled 30 patients with TBI who underwent diffusion tensor imaging and language function evaluation using the Western Aphasia Battery. The CC was divided into five segments (C1-C5) according to its projecting fibers using Hofer’s method, and fractional anisotropy (FA) values were measured using DSI studio software. The FA values of the left arcuate fasciculus and cingulum for language function and executive function, respectively, were also evaluated. Twelve healthy controls were also enrolled to compare the FA values of these tracts.

**Results:** The FA values of the cingulum and left arcuate fasciculus were significantly correlated with all language scores. The FA values of the entire CC were significantly correlated with the fluency, repetition, and aphasia quotient scores. The FA values of the anterior CC segment (C1 and C2) significantly correlated with the aphasia quotient score; C1 with the fluency score; and C2 with the fluency, comprehension, and repetition scores. However, the FA values of the posterior CC (C3-C5) were not significantly correlated with any of the language subset scores.

**Conclusion:** The language function in patients with TBI is correlated with the integrity of the white matter tracts important for language and attention processes. Moreover, disruption of the CC is common after TBI, and the anterior CC segment plays an important role in language impairment after TBI. Therefore, analyzing CC integrity using diffusion tensor imaging can help predict language impairment in patients with TBI.

## Introduction

Traumatic brain injury (TBI) occurs after an external force injury to the brain, such as acceleration/deceleration, concussion, or penetration [1]. The recovery pattern is quite different from that after brain infarction or hemorrhage; thus, appropriate rehabilitation intervention is required. The causes of TBI include falls, vehicle collisions, and violence, and its incidence is currently increasing, especially in young populations; it is one of the major causes of morbidity and mortality [1, 2]. The location and severity of TBI are related to shearing forces during acceleration, deceleration, and rotation of the head during injury; the common locations of diffuse axonal injuries include the cerebral hemisphere gray-white matter interface, subcortical white matter, corpus callosum (CC), basal ganglia, dorsal lateral aspect of brainstem, and cerebellum [3].

The CC is the largest commissural white matter bundle in the brain that links the left and right cerebral hemispheres and is one of the most vulnerable sites after TBI, probably owing to its unique midline location and the high possibility of secondary injury due to elevated intracranial pressure [4]. It is of particular interest as it has a structural role in inter-hemispheric transfer of cognitive function, memory, and motor functions [5]. It can be divided into three parts–the genu, trunk, and splenium. The genu connects the prefrontal cortex; the trunk connects the motor and sensory cortices; and the splenium connects the temporal, parietal, and occipital cortices between the two hemispheres. Many studies have been conducted to evaluate the relationship between the CC integrity and cognitive function [6]. Previous studies have identified a positive correlation between the Mini Mental State Examination (MMSE) score and the integrity of the splenium of the CC, and a positive correlation between working memory performance and CC integrity [7, 8]. While many studies have been conducted to determine the relationship between CC integrity and various cognitive measures, only a few studies have focused on the correlation between CC integrity and language function. Although the arcuate fasciculus (AF) remains the main pathway in language processing, the CC is also suggested to play a role in language function, such as language lateralization between the two hemispheres [9, 10].

In this study, we hypothesized that damage to the CC is associated with impairment of language function in patients with TBI. We investigated the association between CC integrity and language function after TBI using diffusion tensor imaging (DTI).

## Materials and methods

### Subjects

Patients with TBI who were admitted to the Physical Medicine and Rehabilitation department of Korea University Anam Hospital in Seoul, Korea, between January 2016 and December 2019 were enrolled. The inclusion criteria included the following: (1) mild-to-moderate TBI, Glasgow Coma Scale (GCS) score of > 8, and Rancho Los Amigos (RLA) scale score of > 4; (2) brain lesion confirmed on computed tomography (CT) or magnetic resonance imaging (MRI); and (3) available data on MRI DTI and language function evaluation using the Korean version of the Western Aphasia Battery (WAB). The exclusion criteria included the following: (1) severe cognitive, behavioral, or physical impairment impeding language function evaluation; (2) history of stroke or degenerative brain disorders, such as dementia or Parkinson’s disease; and (3) persistent critical medical issues, such as ventilator use, after TBI. Finally, a total of 30 patients were eligible for the study. We also included 12 healthy control subjects as the control group for comparison of DTI parameters with the TBI group.

The study was conducted retrospectively, and the study protocol was approved by the Institutional Review Board (IRB) of Korea University Anam Hospital. (IRB No. 2019AN0399)

### Data acquisition

The following data were collected: age, sex, years of education, occupation, radiologic findings (skull fracture, cerebral contusion, traumatic intracerebral hemorrhage, subdural hemorrhage, or epidural hematoma) observed on CT, injury site (frontal, temporal, parietal, occipital, or left or right), GCS score on admission, RLA scale score, MMSE score for cognitive function, and WAB scores for language function evaluation, including fluency (0-20), comprehension (0-10), repetition (0-10), naming (0-10), and aphasia quotient (AQ). The AQ was calculated using the summation of four subset scores multiplied by 2, with a maximum of 100 points.

### DTI acquisition

DTI was performed using a 3.0 T Prisma MRI scanner (Siemens, Erlangen, Germany). High-resolution, structural T1-weighted images of the entire brain were acquired. DTI data were acquired using a spin echo single-shot planar imaging pulse sequence. The image parameters were as follows: number of diffusion gradient directions = 64, matrix = 112 × 112, field of view = 224 × 224 mm2, voxel size = 2.0 × 2.0 × 2.0 mm3, echo time = 55 ms, repetition time = 6500 ms, b = 1000 s/mm2, slide thickness = 2.0 mm, flip angle = 90°. Correction to head movement was conducted using a rigid body transformation method (rotation and translation, six parameters).

### Reconstruction of the white matter tract using DTI

Reconstruction was performed using DSI Studio (http://dsi-studio.labsolver.org). The termination criteria used for fiber tracking were as follows: fractional anisotropy (FA) value of <0.15 and angle change of >70°. The region of interest was placed on the CC, and it was reconstructed by selecting the region of interest using Hofer’s classification [11]. The regions were as follows: region I–the most anterior segment, containing fibers to the prefrontal region, region II–containing fibers to the premotor and supplementary motor cortical areas, region III– containing fibers to the primary motor cortex, region IV–containing fibers to the primary sensory fibers, and region V–containing fibers to the parietal, temporal, and occipital cortices (Fig 1). These regions were named as C1, C2, C3, C4, and C5, respectively. In addition to the CC, we reconstructed the AF and cingulum, which also influence language function and executive attention, respectively. The FA values of the entire CC and the five CC regions as well as those of the cingulum and AF were obtained.

**Fig. 1.**
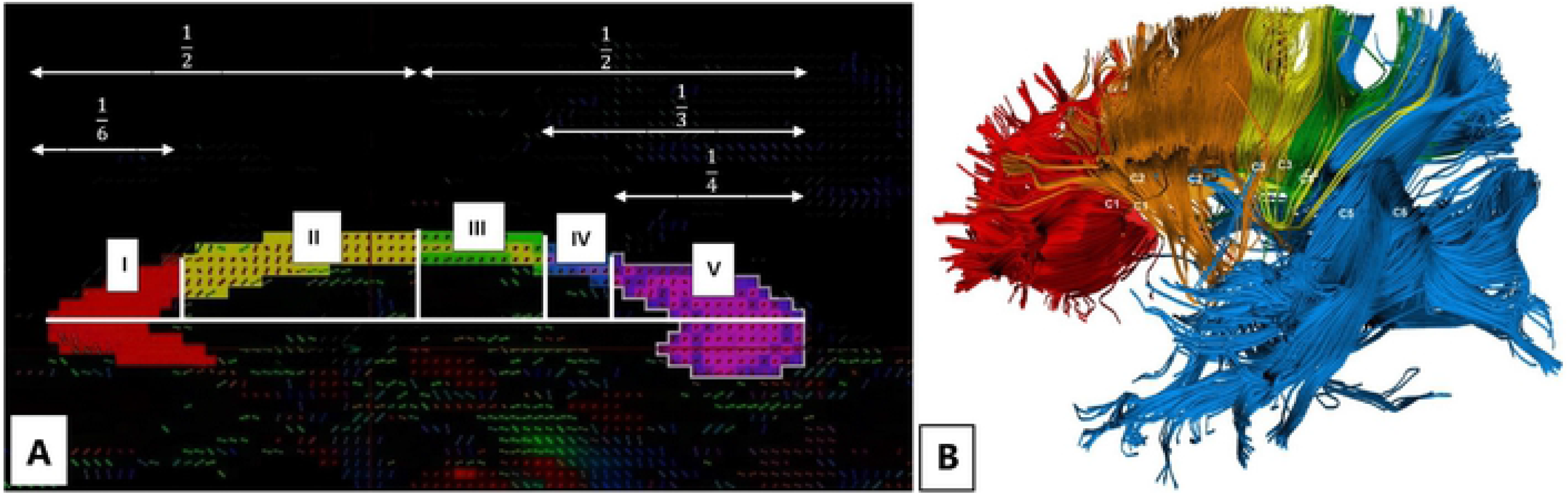
The corpus callosum is divided into five segments according to Hofer’s method (A). The fibers of C1-C5 on diffusion tensor tractography are shown in different colors (B).

### Statistical analyses

Statistical Package for the Social Sciences version 25.0 (IBM, Armonk, NY, USA) was used for the statistical analyses. The Shapiro-Wilk test was used to test the normality of the data. An independent t-test was used to compare the FA values between the TBI and control groups and Pearson’s correlation test was performed for comparisons between the FA values of each tract and language scores evaluated using the WAB. A p-value of <0.05 was considered statistically significant.

## Results

### Characteristics of the subjects

The demographic data of the 30 patients are summarized in Table 1. Among them, 22 were men and 8 were wome; the mean age was 60.7 years. The etiology of TBI was fall or slip-down in 13 patients and motor vehicular accident or other trauma in 17 patients. The average GCS and RLA scale scores were 11.0 and 5.63, respectively, and the mean MMSE score was 12.73 points. The language score evaluated using the WAB was 63.0 for AQ, and the average subset scores were as follows: fluency, 12.58 of 20; comprehension, 6.63 of 10; repetition, 6.77 of 10; and naming, 5.72 of 10. The patients underwent DTI in the subacute phase.

**Table 1.** Demographic and clinical characteristics of the patients with traumatic brain injury (N=30).

| **Variables** | **Mean (Standard deviation)** |
| --- | --- |
| **Age (years)** | 60.77 (17.94) |
| **Sex (male/ female)** | 22 / 8 |
| **Education (years)** | 10.83 (3.10) |
| **Cause of TBI (fall/other)** | 13 / 17 |
| **Glasgow Coma Scale score (3-15)** | 11.0 (4.30) |
| **Rancho Los Amigos scale score(1-8)** | 5.63 (0.96) |
| **MMSE score** | 12.73 (8.37) |
| **Language function (WAB)** |  |
| **Fluency (20)** | 12.58 (4.99) |
| **Comprehension (10)** | 6.63 (3.02) |
| **Repetition (10)** | 6.77 (3.37) |
| **Naming (10)** | 5.72 (3.24) |
| **Aphasia quotient (100)** | 63.0 (27.06) |

TBI: traumatic brain injury, MMSE: Mini-Mental State Examination, WAB: Western Aphasia Battery

### Comparison of the FA values between the TBI and control groups

The control group that underwent MRI DTI consisted of 12 healthy participants (1 man and 11 women, mean age 42.58±15.92 years). The FA values of the entire CC, C1-C5, AF, and cingulum were significantly lower in the TBI group than in the control group. (Table 2)

**Table 2.**
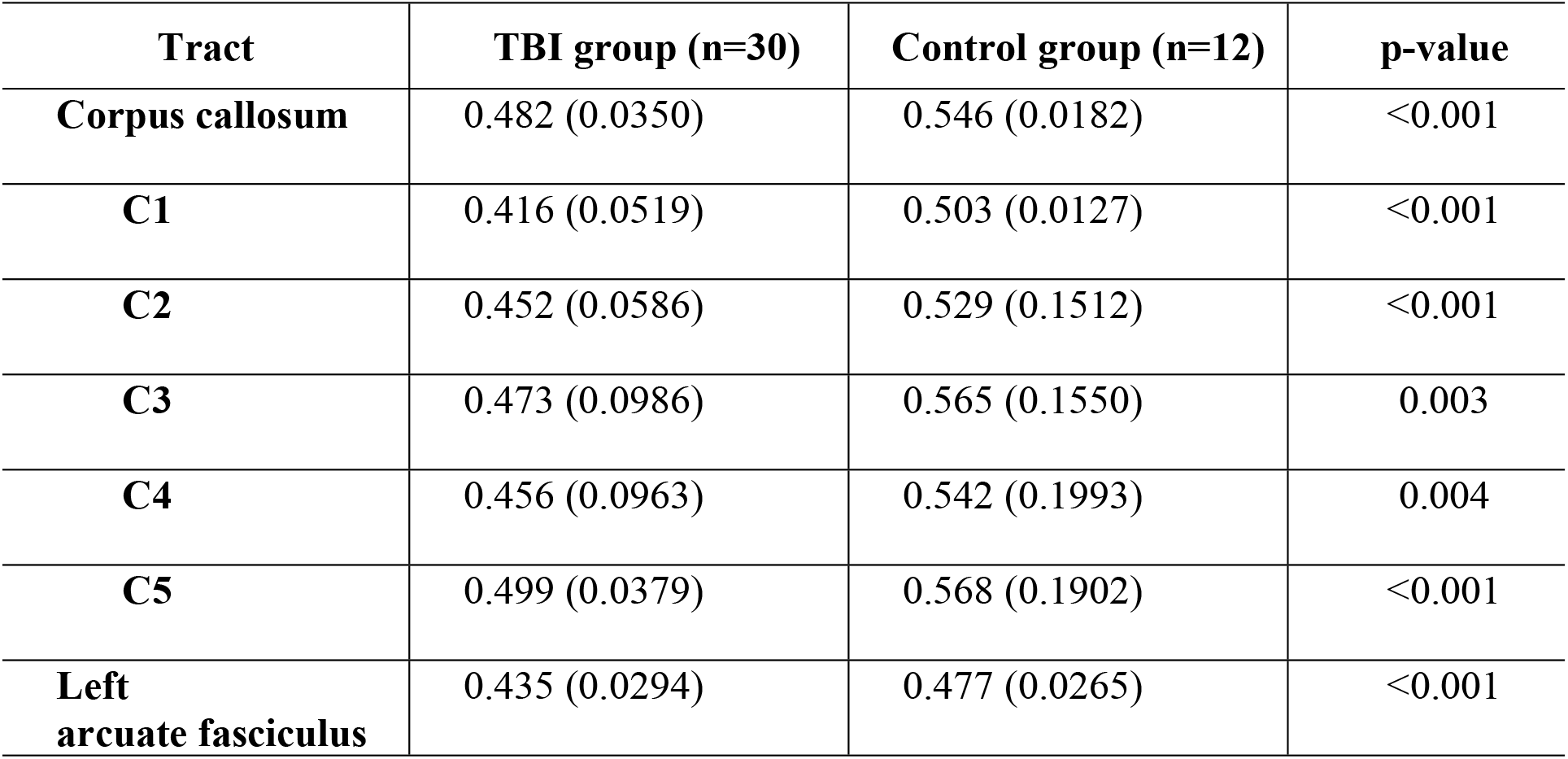
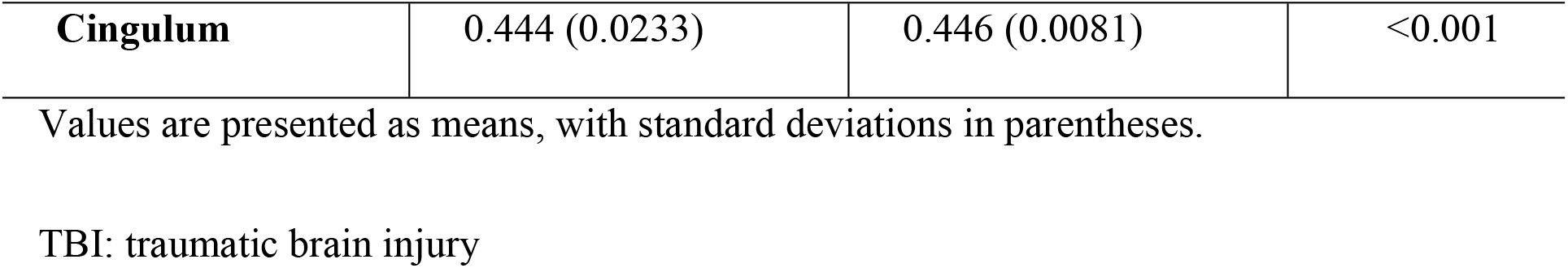
Comparison of the FA values for the corpus callosum, arcuate fasciculus, and cingulum between the TBI and control groups.

### Correlation between the FA values of the CC and language scores

Pearson’s correlation test was used to identify the relationship between the FA values of each tract and the WAB scores in the TBI group. The correlation analysis results between the FA values of each tract and the WAB scores are provided in Table 3. The FA values of the left AF and cingulum showed a significant correlation with all the language subset scores. Compared with the values of the AF and cingulum, the FA values of the entire CC were significantly correlated with the fluency, repetition, and AQ scores. The FA values of the anterior segment of the CC (C1 and C2) were significantly correlated with the overall language function presented by the AQ score. The FA value of C1 was significantly correlated with the fluency score and that of C2 was significantly correlated with the fluency, comprehension, and repetition scores. However, the FA values of the posterior segment of the CC (C3-C5) were not significantly correlated with any of the language subset scores.

**Table 3.**
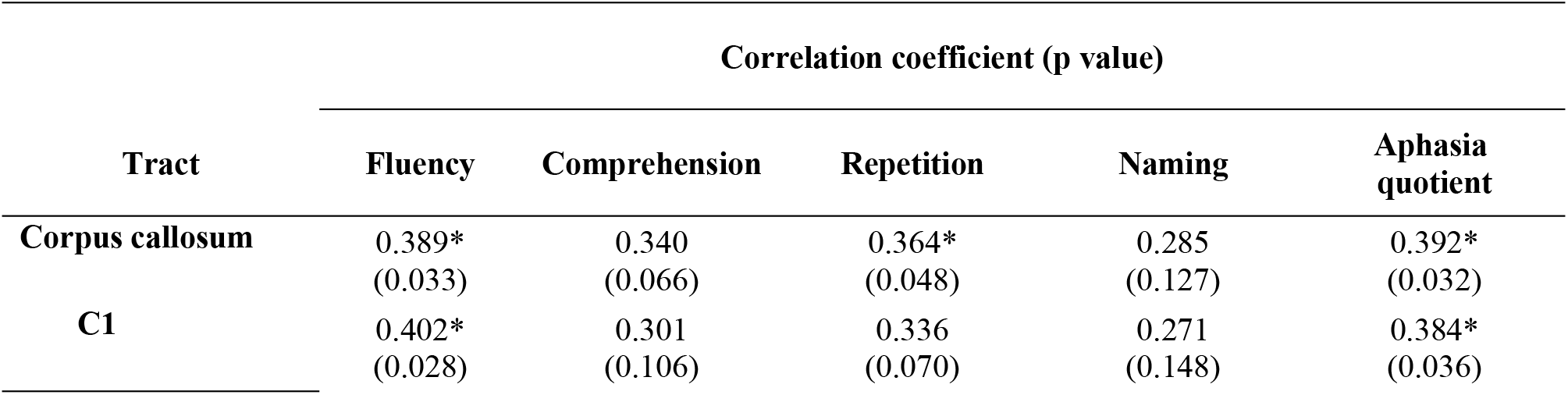
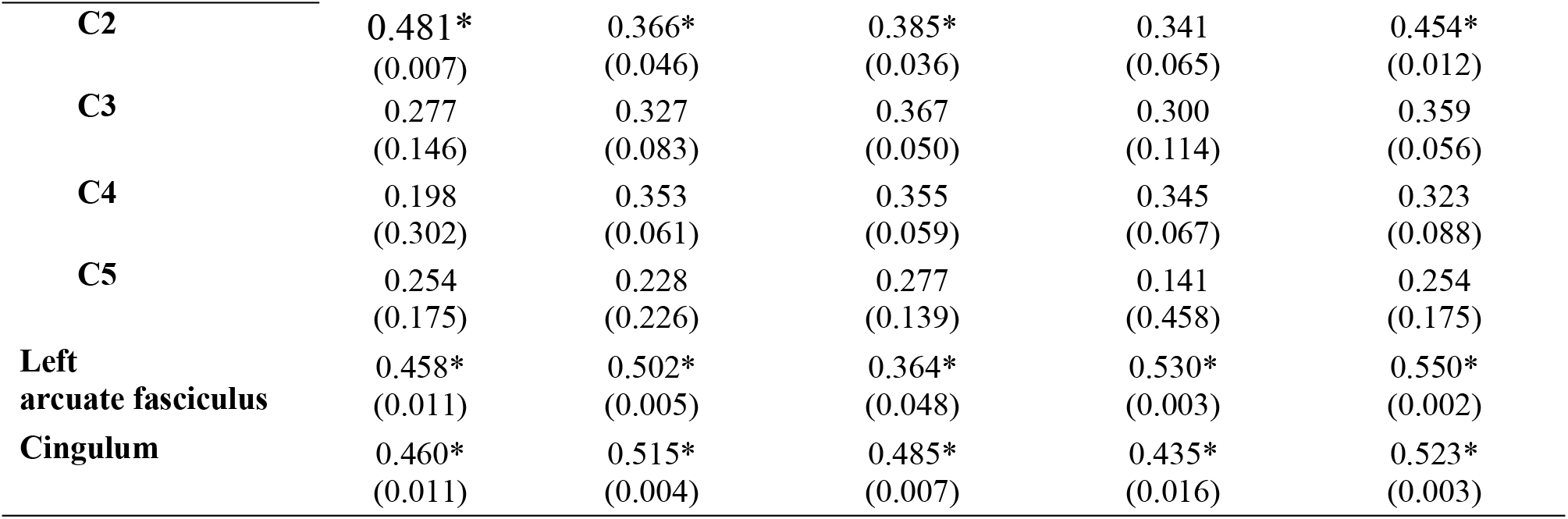
Correlation coefficient of the FA values in relation to the language scores of patients with traumatic brain injury.

## Discussion

This study aimed to identify the correlation between CC integrity and language function in patients with TBI. Our analysis showed that the overall integrity of the white matter tracts of the CC, AF, and cingulum was significantly affected after TBI. The FA values of the entire CC and its segments (C1-C5) were considerably lower in the TBI group than in the control group. Only FA values of the anterior segment of the CC (C1 and C2) showed a significant correlation with language function.

DTI has been widely used in studies on white matter tract injuries among patients with TBI because it can detect minor abnormalities that conventional CT or MRI cannot [12]. In TBI, consistent decrease in anisotropic diffusion has been reported in the chronic phase after TBI. However, considerable debate remains if this occurs in the acute and sub-acute phases [13]. Although the CC is one of the predilection sites of traumatic axonal injury, it is believed to be unevenly affected after trauma [4, 5]. In this study, the FA values of the AF, cingulum, and CC were significantly lower in the TBI group than in the control group. Further, the FA values of all segments of the CC consistently decreased, suggesting vulnerability of the CC after brain injury.

The majority of patients with TBI do not demonstrate classic aphasia symptoms [14], and executive function deficit does not solely explain this [14, 15]. There is no defined role of the CC in language processing; however, some studies have suggested its role in language lateralization and non-literal language processing which are important in social communication [10, 16]. Some studies have reported cases of aphasia after CC injury; however, the precise mechanism has not been illustrated [17–19]. One proposed theory is that the equilibrium between the bilateral hemispheres changes after trauma; damage in the left hemisphere language center results in diaschisis between the bilateral cortices and the excitability in the right hemisphere language mirror area increases. Recovery requires reestablishment of excitation balance between the bilateral hemispheres and CC, the largest pathway that connects the two hemispheres, plays a critical role in the transcallosal diaschisis in the language area so that aphasia recovery occurs [20, 21]. Previous studies have suggested that the anterior segment of the CC shows DTI abnormalities after trauma, making it more associated with poor prognosis [5, 7, 22, 23]. One study suggested a different mechanism underlying injury in the genu and splenium: FA reduction in the genu was accompanied by apparent diffusion coefficient (ADC) increase, whereas FA reduction in the splenium occurred in the absence of ADC change. This indicates that splenium damage is most likely to be irreversible, and genu damage is more reversible [5].

In this study, the FA values of the CC and all the segments overtly decreased in the TBI group compared to the control group. We believe that despite the fact that the integrity of all CC segments was disrupted after trauma, different segments contribute to language impairment. In our analysis, the FA values of the anterior CC segment (C1 and C2) showed a significant correlation with the overall language function (AQ score). Further, the FA value of C1 was associated with the fluency score; and that of C2 was related to the fluency, comprehension, and repetition scores. However, the FA values of the posterior segment of the CC showed no significant correlation with any of the language subset scores. Our results suggest that the FA values of the anterior segment of the CC have a greater relationship with language impairment after TBI, especially the fluency score among all the language subset scores. One previous study has revealed that patients who had their anterior segment of the CC (genu and/or body) surgically removed demonstrated deficits in motor behavior and transfer of somesthetic information and that the anterior portion of the genu is associated with asynchronous motor coordination and planning [24, 25]. In another study, 4 out of 36 patients with callosal infarcts had decreased verbal fluency [20]. Further, the CC undergoes structural and functional reorganization of the language network after injury, which accounts for the improvement of spontaneous speech and repetition [18]. Thus, previous studies have observed that most patients with callosal injury demonstrats verbal language impairments. Taken together, it is reasonable to expect that because the anterior segment of the CC is most likely to be disrupted after trauma, this part plays an important role in language function after TBI.

Some limitations of this study include its design; this study was a single institutional, retrospective study. Further, the number of patients who participated in the study was small because many patients with TBI did not undergo DTI and language evaluation owing to severe cognitive behavioral symptoms and comorbidities. A large clinical trial should be conducted in the future to better define the relationship between CC lesions and language function. A longitudinal study that would demonstrate the long-term effects of CC integrity disruption on language function should also be performed to further understand the underlying mechanisms.

## Conclusion

In conclusion, the language function in patients with TBI is correlated with the integrity of the white matter tracts important for language and attention processes. Moreover, disruption of the CC is common after TBI, and the anterior segment of the CC plays an important role in language impairment after TBI. Therefore, analyzing CC integrity using DTI can help predict language impairment in patients with TBI.

## Acknowledgements

We would like to thank Editage (www.editage.co.kr) for English language editing.

